# *E. coli* Nissle 1917 occupies previously undocumented host niches in the insect-parasitic nematode *Steinernema hermaphroditum*

**DOI:** 10.1101/2025.07.03.663106

**Authors:** Victoria Chen, John P. Marken, Richard M. Murray, Mengyi Cao

## Abstract

*Steinernema* species are soil-dwelling and insect-parasitic nematodes that associate with symbiotic *Xenorhabdus* bacteria. During the infective juvenile (IJ) stage*, Steinernema* nematodes package species-specific *Xenorhabdus* bacteria in the anterior intestinal pockets. The nematodes can survive in the soil for months as they seek insect prey. The mechanisms of how these nematodes associate with environmental microbes other than their *Xenorhabdus* symbionts is barely known. Here, we report a new mechanism of *E. coli* Nissle (EcN) association with the nematode *Steinernema hermaphroditum.* We show that EcN cells are enclosed and lysed in at least four pairs of coelomocytes, suggesting these immune cells respond to bacterial invasion. During the IJ stage of nematode development, EcN cells localize to posterior intestinal vacuoles and enter the inter-cuticular space, where they proliferate, aggregate, then lyse. EcN cells expressed proteins in the cell lysates were maintained in the nematode cuticle over eight weeks in non-sterile soil. These observations suggest sequential steps of EcN colonization in the host nematodes involving an immune response that is distinctive from interactions with mutualistic symbiont. Our work establishes a novel framework of nematode-bacteria interaction with potential applications in environmental bioengineering.

## Introduction

Animal tissues create complex, compartmentalized spaces that provide abiotic and biotic environments capable of selecting and maintaining mutualistic microorganisms while antagonizing pathogenic ones. Reciprocally, microbes develop within these host-derived environments, shaping the identity of the niches they occupy. This process of microbial modulation of the local environment, known as niche construction, can drive long-term coevolution. In contrast, pathogens can exploit unoccupied niches, leading to opportunistic infections and niche deconstruction (Baquero *et al*., 2021). Identifying and characterizing novel niches within animal hosts, as well as the dynamic interactions of microbes within these microenvironments, are crucial for understanding the mechanisms and consequences of host-microbe crosstalk. Binary symbioses, involving a single core symbiont and a single animal species, provide efficient models for revealing the temporal and spatial dynamics of symbiont colonization within specific host cells and tissues. The niche-specific interactions observed in diverse binary symbiosis models are critical for host development and ecosystem health (Nyholm and McFall-Ngai, 1998; Stilwell *et al*., 2018; Maire *et al*., 2020; Tivey *et al*., 2022). While binary symbiosis often involves colonization by a single microbial species within a specialized organ, additional genera or species can sometimes associate with distinct tissues, maintaining spatial separation from primary symbionts (Dale and Moran, 2006; Nyholm and McFall-Ngai, 2021). The relatively simple composition of these natural microbiomes makes such animal models advantageous for studying multiple types of host-microbe interactions simultaneously.

The entomopathogenic (EPN, insect-parasitic) nematode *Steinernema spp* maintains a stable association with mutualistic *Xenorhabdus* bacteria in a binary symbiosis. Each generation of nematodes acquires their symbiotic bacteria from the external environment (semi-horizontal) while a maternal factor can influence the bacterial colonization of the next generation (semi-vertical). The intestine of *Steinernema* nematodes contains compartmentalized niches where multiple microenvironments enable the attachment, persistence, and proliferation of *Xenorhabdus* bacteria (Chaston *et al*., 2013) (Fig. 1A and Fig. 1B). *Xenorhabdus* bacteria first attach to the anterior intestinal caecum (AIC) in juveniles, while nematodes feed on bacteria, molt through four juvenile stages (J1-J4) and become adults to reproduce. Signals from overcrowding and lack of food source induce the J2 stage of juveniles to molt into IJs through an alternative developmental pathway. During IJ development, the symbiotic bacteria first localize to the pharyngeal intestinal valves (PIV) in the pre-IJs. These narrow pouches restrict bacterial colonization to a few cells per animal. During the infective juvenile (IJ) stage, symbiotic bacteria migrate into the anterior intestinal pocket, termed the receptacle (Fig. 1B). The multiple intestinal tissues involved in the colonization events, including AIC, PIV, and the receptacle, are proposed to provide specific nutrients, signaling molecules, and physical restrictions to select the specific species or strain of *Xenorhabdus* bacteria.

**Figure 1:**
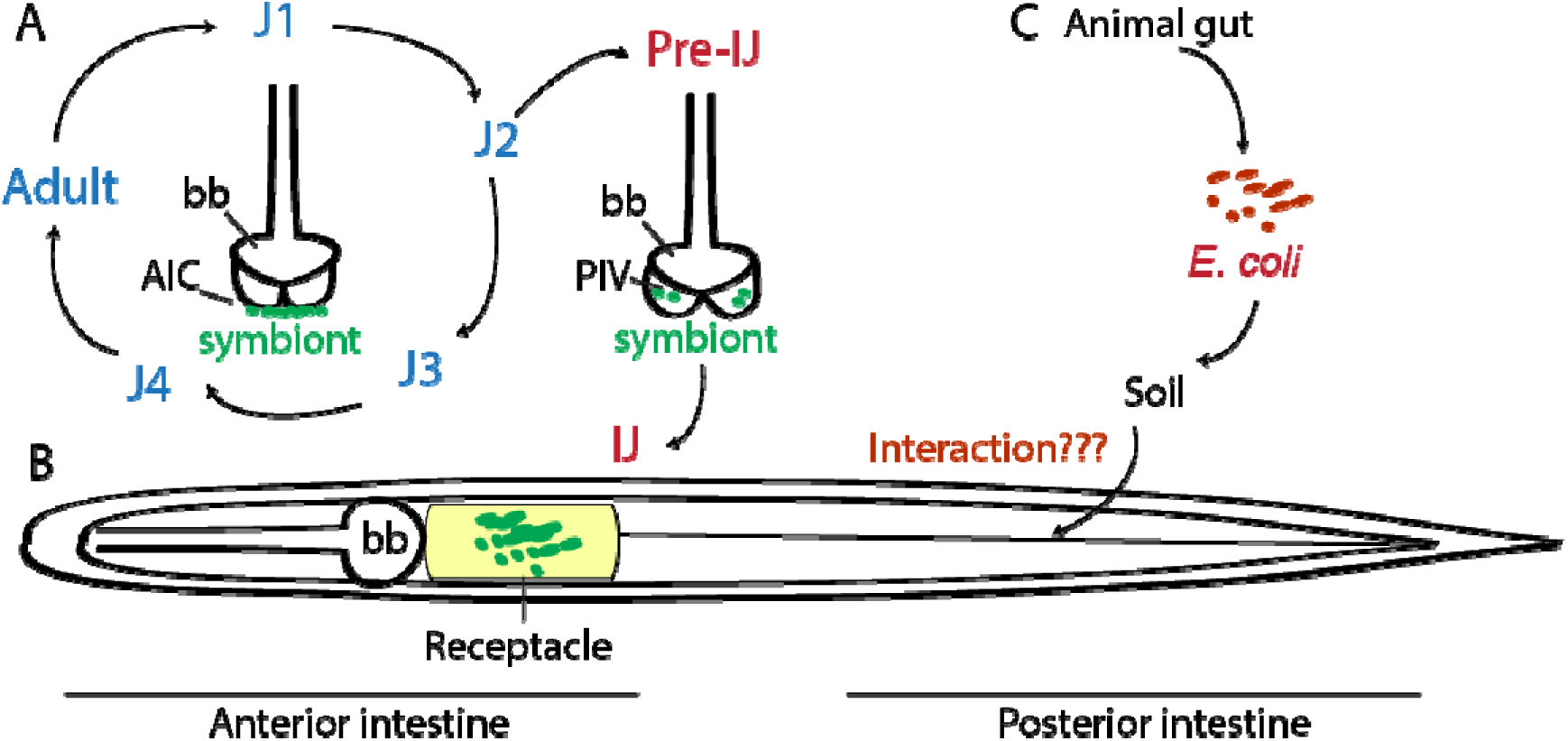
*Steinernema hermaphroditum* intestine provides specific niches for host-microbe interactions. (A). *S. hermaphroditum* life cycle and symbiont colonization in specific tissues at the anterior intestine. (B). *S. hermaphroditum* IJ that has the potential to interact with both mutualistic and pathogenic microbes. (C). Biphasic lifestyle of *E. coli* that oscillates between associating with mammalian gut and persisting in the soil, which provides an opportunity to interact with soil-dwelling *Steinernema* nematodes. Bb: basal bulb; AIC: anterior intestinal caecum; PIV: pharyngeal intestinal valve.

*Steinernema* are soil-dwelling nematodes that also encounter other species of microbes from their natural environment. Bacteria from multiple genera, termed the ‘pathobiome’, are found to frequently associate with certain *Steinernema* spp and potentially contribute to entomopathogenic (insect-killing) activities (Ogier *et al*., 2020). Despite its generalization of pathobiomes among multiple *Steinernema* species, the frequency, localization, and molecular mechanisms of pathobiome association is unknown. Little else is known about other environmental microbes -- either pathogenic, mutualistic, or commensal -- that interact with EPNs. This knowledge gap limits the full application of *Steinernema* as a symbiosis model and its applications in bioengineering. Therefore, understanding the spatial and temporal interactions of pathogens and commensals in *Steinernema* provides a foundational framework for novel modes of symbiotic interactions.

*Escherichia coli* is the most established bacterial species for molecular biology and is a leading candidate for bioengineering applications across diverse niches, including biosensing in complex microenvironments (Del Valle *et al*., 2021) and modifying gene expression in multicellular organisms (Gao and Sun, 2021). *E. coli* is commonly associated with the animal intestinal tract (host-dependent) and circulates through the soil and water environment (host-independent) via a biphasic lifestyle (Fig. 1C). Certain *E. coli* strains can persist in soil environments, where they influence indigenous microflora encountered by insects and soil-dwelling nematodes (Elsas *et al*., 2010). The *E. coli* Nissle 1917 (EcN) strain, originally isolated from a soldier who survived a *Shigella* endemic, is known to interact with the mammalian intestinal epithelium and mucosal immune system in a non-pathogenic manner, antagonize pathogenic enterobacteria, and confer probiotic benefits in humans (Sonnenborn, 2016). In model nematode *Caenorhabditis elegans* adults, EcN as food source was observed to show both probiotic and neurodegenerative effects (Kim and Moon, 2019; Redweik and Xue, 2025) . Engineered EcN were used to carry biosensors that monitor the intestinal environment of the animal (Rutter *et al*., 2019). However, the interactions of EcN with nematode species beyond *C. elegans* is not explored. Understanding the interactions between EcN and *Steinernema* are therefore useful for establishing a foundation for potential synthetic biology solutions to challenges in host-microbe interactions and application in the soil environment (Jones *et al*., 2023) .

In this study, we investigated the interactions between *E. coli* Nissle and the entomopathogenic nematode *Steinernema hermaphroditum*. We discovered that EcN localizes within previously undocumented cell and tissue types in *S. hermaphroditum*, including immune cells (coelomocytes), posterior intestinal tissues, and inter-cuticular spaces. These snapshots of EcN localization in nematodes suggest a sequential process of cell- and tissue-specific microbe-host interactions synchronized with infective juvenile (IJ) development. Notably, EcN colonization occurs in spatially distinct tissues from those occupied by *Xenorhabdus*, the nematode’s native symbiont. Our findings established *E. coli* Nissle–*S. hermaphroditum* interactions as a model to monitor niche-specific microbial interactions. As an emerging genetic model organism, *S. hermaphroditum* offers a promising delivery system for engineered biosensors due to its transparency, short life cycle in vitro, and bacterivorous nature (Bracho *et al*., 2016; Hwang *et al*., 2017; Rutter *et al*., 2019).

## Experimental Procedures

### Construction of fluorescent *E. coli* Nissle strains with mScarlet-I

Genetic constructs for constitutive expression of red fluorescent protein (mScarlet-I) was assembled using the 3G assembly method (Halleran *et al*., 2018) with standard parts from the MoClo modular cloning toolkit (Iverson *et al*., 2016). The resulting constructs were cloned into the “KL” genome integration vector from the pOSIP one-step genome integration system (St-Pierre *et al*., 2013), which facilitates integration at the lambda integrase attachment site in the *E. coli* genome. Plasmids were transformed into chemically competent *E. coli* strains (Nissle 1917, CSH50 (Sokurenko *et al*., 1992), and BW25113 (Grenier *et al*., 2014)) using a standard heat shock protocol. Briefly, 50 µL of competent cells were mixed with 100 ng of plasmid DNA, incubated on ice for 30 minutes, heat shocked at 42 °C for 45 seconds, and then placed back on ice for 2 minutes. Cells were allowed to recover in 950 µL of SOC medium at 37 °C with shaking for 1 hour before being plated on LB agar plates containing kanamycin (50 µg/mL) to select for successful integrants. Plates were incubated at 37 °C overnight. Resulting transformants were screened via colony PCR on the integration locus using the standard screening primers from (St-Pierre *et al*., 2013) and Sanger sequenced to validate the successful integration.

### Liver-kidney agar plates

The recipe for Liver-kidney agar was adapted from (Flores-Lara *et al*., 2007): 50 g of pork liver and 50 g of pork kidney were blended in 250 mL distilled water until smooth. 2.5 g of NaCl, 7.5 g agar, 250 mL of water, and the liver-kidney solution was transferred to a flask, then autoclaved and cooled in a 55°C water bath. To avoid contamination, the liver-kidney agar was supplemented with kanamycin (50 μg/mL) and poured into 60 x 15mm Petri plates under a fume hood. The agar was continuously swirled before pouring to avoid pouring any larger chunks.

### *S. hermaphroditum* axenic egg extraction and colonization assay

Conventional *S. hermaphroditum* nematodes were maintained by propagating through *Galleria mellonela* 5^th^ instar larvae (Cao *et al*., 2021). The method for axenic egg extraction for *S. hermaphroditum* was adapted from (Murfin *et al*., 2012). Liquid cultures of symbiotic bacterium *X. griffiniae* (HGB2511) were grown overnight in LB media that were kept in the dark and added to lipid agar plates (Vivas and Goodrich-Blair, 2001; Cao *et al*., 2021). Plates were cultured for 48 hours to form a bacterial lawn. To prepare axenic (germ-free) eggs, conventional IJs (exclusively propagated through insects) were added to the lawns of *X. griffiniae* (HGB2511) and incubated for 2-3 days to grow to gravid adults. Nematodes were washed off the bacterial lawn with water and then transferred to 50 mL conical tubes (Falcon, Corning, NY) and centrifuged at 3000 rpm for 2 minutes. The supernatant was removed using a vacuum, and 2 mL of a lysis solution (2.4 mL 8% bleach, 500 µL 5M KOH, 7.1 mL water) was added to lyse the gravid adult hermaphrodites. Nematodes were incubated in the lysis solution for 5 minutes at room temperature and mixed several times per minute by inverting. The solution was then centrifuged at 3000 rpm for two minutes, and the lysis solution was removed using a vacuum. Nematode eggs were washed thrice with 10 mL of LB medium. Their density was assessed by viewing the number of IJs in 2 µL spots under a stereoscope.

For colonization assays, approximately 500 eggs were added to the prepared liver-kidney or lipid agar plates, with or without bacterial lawn of *X. griffiniae* expressing GFP (Thomas *et al*., 2024) or *E. coli* expressing mScarlet-I and incubated at 25℃ for 1 week. Plates were then moved to 100 x 25mm Petri dishes filled with 10-15 mL of water to trap infective juveniles. Three biological replicates of each colonization condition were screened to identify colonization phenotypes based on the presence of EcN in different tissues. Colonization phenotypes and frequencies were assessed using fluorescence microscopy (See Microscopy and Image Acquisition for details).

### Calculation of CFU per IJ

The CFU per IJ assessment was adapted from (Murfin *et al*., 2019). IJ suspension from water-trap was added to a sterile 1.5 mL microfuge tubes (Eppendorf, Germany) and centrifuged at 634 g for two minutes to concentrate the nematodes. The supernatant was removed using a vacuum, and 1 mL of a 1% bleach solution was added to the pellet, which was incubated for 2 min at room temperature to surface sterilize IJs. IJs were washed thrice with and resuspended in 1 mL of LB medium, and their concentration was assessed by viewing 2 µL spots under a stereoscope. 200 IJs were then resuspended in 250 µL of LB broth. IJs were homogenized for 2 minutes using a hand-held motor-driven grinder and Kontes polypropylene microtube pellet pestle. Serial dilutions of the homogenate were then plated on LB agar plates with 50 μg/mL kanamycin.

### Soil Challenge of IJ colonization

IJ suspensions were transferred to sterile 15 mL conical tubes and centrifuged at 3000 rpm for 2 minutes to form a pellet. IJs were then resuspended in 10 mL of M9 buffer (Stiernagle, 2006) and split into two 50 mL culture flasks, 5 mL each. The concentrations of IJs in each flask were assessed under a stereoscope and recorded. For each plate, a 1:5 dilution of M9-soaked soil (5 g Miracle-Gro Potting Mix in 45 mL of M9 incubated at room temperature overnight, approximately 8000 CFU/µL) was filtered through a Whatman paper and added to one of the flasks to simulate microbial challenge. Weekly, the colonization frequency for each flask was measured via fluorescence microscopy by counting the number of colonized and uncolonized worms. 200 IJs were removed from the flask, paralyzed with 2 mM levamisole in 10 µL of sterile water, and imaged on 5% agar pads.

### Microscopy and Image Acquisition

IJ nematodes were concentrated from water-traps by centrifugation at 1761 g for one minute and immobilized by addition of levamisole to a final concentration of 5 mM. The nematodes were mounted onto 5% agarose in water and covered with a cover glass (Cao *et al*., 2021) . Photomicrographs and videos were acquired using Zeiss Imager Z2 microscope equipped with Apotome 2 and Axiocam 506 mono using Zen 2 Blue software.

## Results

### *E. coli* Nissle cells interact with coelomocytes of *S. hermaphroditum* IJ

Using fluorescence microscopy, we observed that mScarlet-I-expressing *E. coli* Nissle cells localize to at least four ‘spots’ on the lateral sides of *S. hermaphroditum* IJs, with two spots on the posterior dorsal side and two on the anterior ventral side (approximately 5-10 μm in diameter) (Fig. 2A and 2B). Z-stack imaging for these four ‘spots’ showed that each consists of a pair of large ovoid cells (Fig. 2D, Video S1). Within each cell, the cytosol, containing lysates from fluorescent bacteria, surrounds a dark center which features the nuclease (Fig. 2D). Based on the morphology and localization of these cells, we identify them as the coelomocytes, a cell type known to serve defense and immune functions in invertebrates including some nematode species (Engelmann *et al*., 2005; Tahseen, 2009; Smith *et al*., 2010). Nematode coelomocytes are known to have diverse morphologies, sizes, and numbers. The morphology of coelomocytes ranges from oval-shaped to star-shaped; the sizes of these cells correlate with the size of the animals (from less than 1 mm to a few meters long). They are proposed to participate in the up-take and scavenging of proteins circulating in the pseudocoelomic space. Whether or not the coelomocytes could engulf bacterial cells depends on the species of nematode (Tahseen, 2009). For instance, *C. elegans* coelomocytes were found to only have endocytosed toxins, such as nematode-expressed green fluorescent proteins that circulate in the body cavity, but not pathogens (Fares and Greenwald, 2001). In contrast, the mammalian parasitic nematode *Ascaris suum* was observed to phagocytose living bacterial cells in their coelomocytes (Tahseen, 2009). Neither the morphology nor the function of *Steinernema* coelomocytes has been previously documented. As expected, we observed that *S. hermaphroditum* coelomocytes are oval-shaped, like other *Rhabditis* nematodes; rod-shaped bacterial cells were visible in the coelomocytes (Fig. 2C, 2E, and 2F) consistent with the model that bacteria are trapped and lysed in the lysosomes. These observations suggest the phagocytotic and endocytic functions of *Steinernema* coelomocytes directly towards invading *E. coli* Nissle which causes an active immune response in the host nematode.

**Figure 2:**
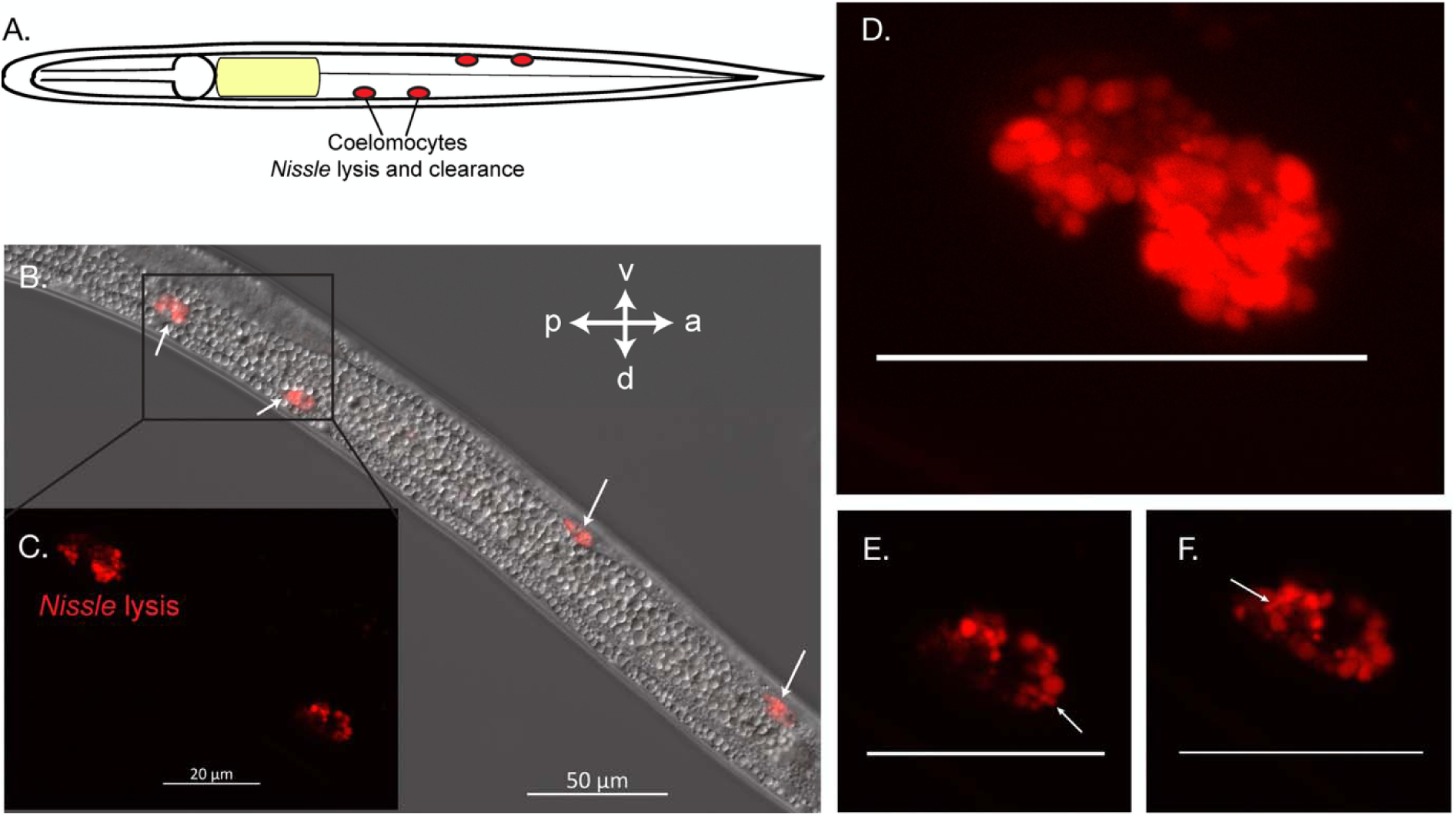
*E. coli* Nissle localizes in the *Steinernema hermaphroditum* IJ coelomocytes. (A). A schematic diagram showing *S. hermaphroditum* IJs has at least four pairs of coelomocytes. (B) and (C). A representative IJ showing *E. coli* Nissle localized and lysed within coelomocytes. (D). Each ovoid coelomocyte structure contains two cells. (E) and (F). Rod-shaped bacterial cells are observed inside of the coelomocyte cytosol. Scale bars in panel D, E, and F equals to 20 µm. (Also see Video S1).

### *E. coli* Nissle cells localize and proliferate in the IJ intestine

As typical of bacterivorous nematodes, the intestine of *Steinernema* serves as the largest organ facilitating microbial interactions where bacteria can be engulfed, digested, or colonized (McGhee, 2007) . Prior to IJ development, EcN cells are engulfed and digested in the intestinal lumen of *S. hermaphroditum* juveniles (Fig. S1). When the intestinal lumen collapses and closes during IJ development, we observe EcN cells attaching to the posterior end of the IJ intestine within multiple small ‘pockets’. We term these structures, which were not previously identified, as ‘intestinal vacuoles’ (Fig. 3A and 3B). Within the same IJ, EcN cells localize and proliferate in one or more intestinal vacuoles (Fig. 3B), showing their role as reservoirs for invading microbes. Individual bacterial cells were also observed to ‘escape’ from the vacuoles, pass through the anus, and eventually enter the space in between the internal cuticle and a layer of growing exterior cuticle (Fig 3B, panel c). This process may be caused by either bacterial motility (such as swimming), or by the movement of the nematode intestine during IJ development as it collapses and squeezes out the bacteria accumulated inside the intestinal lumen. Intestinal colonization was not observed for two other tested laboratory strains of *E. coli*: strain BW25113 and strain CSH50, suggesting that *E. coli* Nissle contains genetic factors that allow colonization not found in these other laboratory strains (Fig. 3C). Intestinal colonization was specific to liver-kidney agar condition lacking *X. griffiniae* symbiotic bacteria (Fig. 3C), therefore, nutritional profile of liver-kidney agar and the specific features of the *E. coli* Nissle strain are essential. The addition of the *X. griffiniae* symbiont was also shown to inhibit intestinal colonization of EcN (Fig. 3C), possibly due to its ability to antagonize *E. coli* growth by producing antimicrobial molecules (Dreyer *et al*., 2018). Overall, EcN is likely a mild and opportunistic pathogen of *S. hermaphroditum,* and some EcN cells can gain resistance to the host immune response by invading and persisting in the IJ intestine.

**Figure 3:**
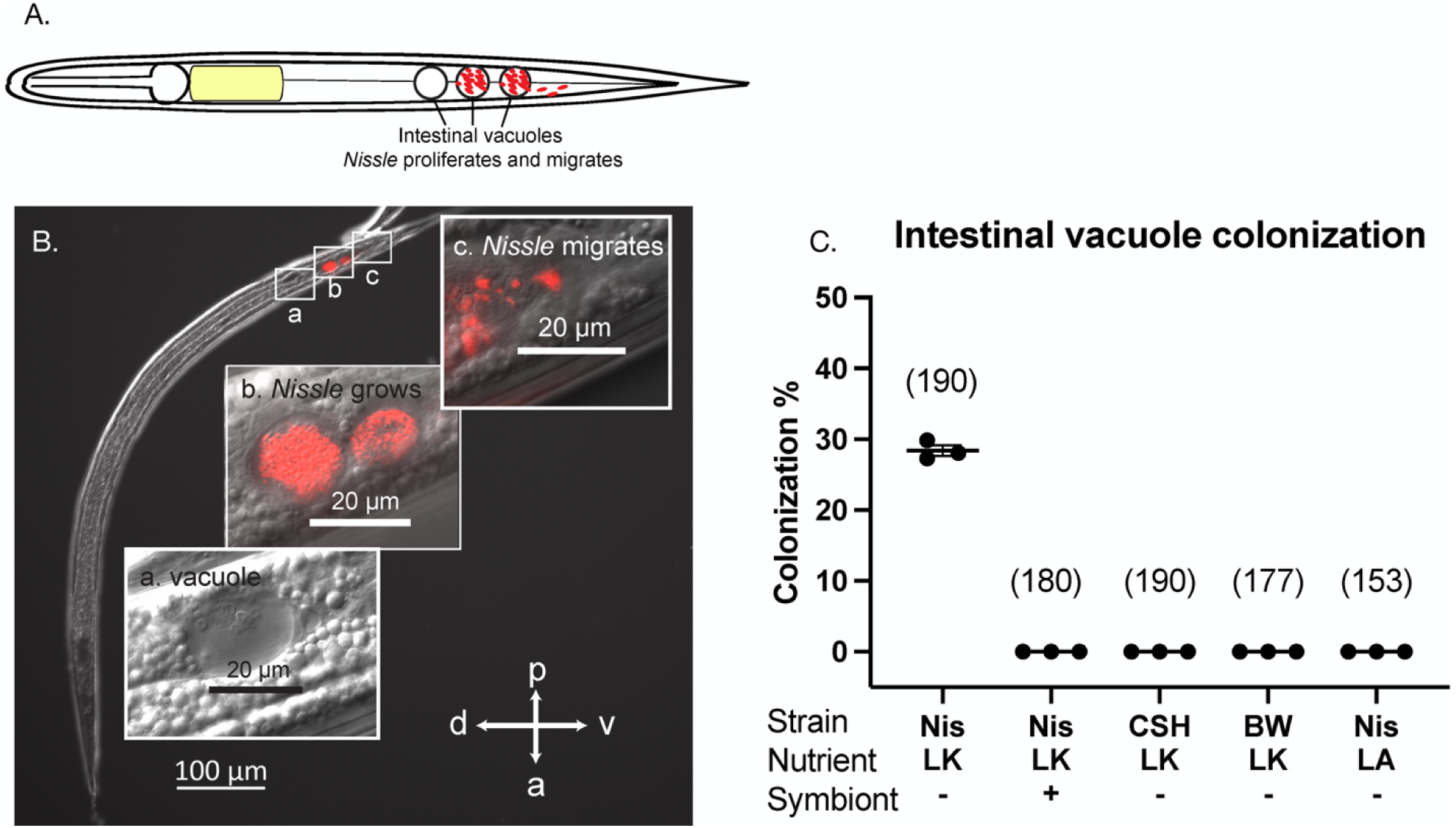
*E. coli* Nissle localizes in the *S. hermaphroditum* intestinal lumen and vacuoles. (A). A schematic diagram showing *E. coli* Nissle invading nematode intestine and proliferating in the intestinal vacuoles. (B). A representative IJ showing intestinal colonization of bacteria: an empty intestinal vacuole containing cell or organelle debris-looking substance (a); bacterial colonization in the IJ intestinal vacuoles (b); bacterial cells escape from the posterior intestinal opening (c). (C). Quantification of intestinal colonization frequency of *E. coli* strains: Nissle (Nis), *E. coli* CSH50 (CSH), BW25113 (BW) under nutritional conditions of either liver-kidney (LK) or lipid agar (LA). Treatments were either with (+) or without (-) symbiont *X. griffiniae.* Numbers in the parenthesis indicate the number of animals screened in each treatment.

### *E. coli* Nissle cells aggregate and migrate in the IJ inter-cuticular space

*S. hermaphroditum* IJs can have up to two cuticle layers: an inner collagenous cuticle similar to that seen in other developmental stages (Wolkow and Hall, 2011), and sometimes an outer cuticle that is known to protect the animal from pathogenic microbes (Timper and Kaya, 1989). We observed aggregates of bacterial cells trapped in the posterior end of the inter-cuticular space (Fig. 4A and 4B; Fig. S3). Cuticular association of EcN is either exclusively localized to the posterior side or co-localized at both the posterior and anterior sides (Fig. S3). No IJ was only associated at the anterior cuticle without posterior colonization, confirming the directionality of colonization from the posterior to the anterior; thus, this event must follow intestinal colonization. Like colonization of the posterior intestinal vacuoles, inter-cuticular colonization was also specific to growth on liver-kidney agar without the presence of any *X. griffiniae* symbionts, and was not observed for the laboratory *E. coli* strains BW25113 or CSH50 (Fig. 4C). This is consistent with our hypothesis that these two colonization phenotypes are correlated (Fig. 3C; Fig. 4C). Along this line of evidence, we also observed the co-occurrence of EcN cells during intestinal and cuticular colonization (Fig. S3). We attempted to assess the CFU per cuticle by surface-sterilizing and grinding IJs, and growing EcN cells on the LB agar plates. However, bleaching process removed the outer cuticle and killed the EcN cells in the inter-cuticular space, causing nearly 0 CFU per IJ.

**Figure 4:**
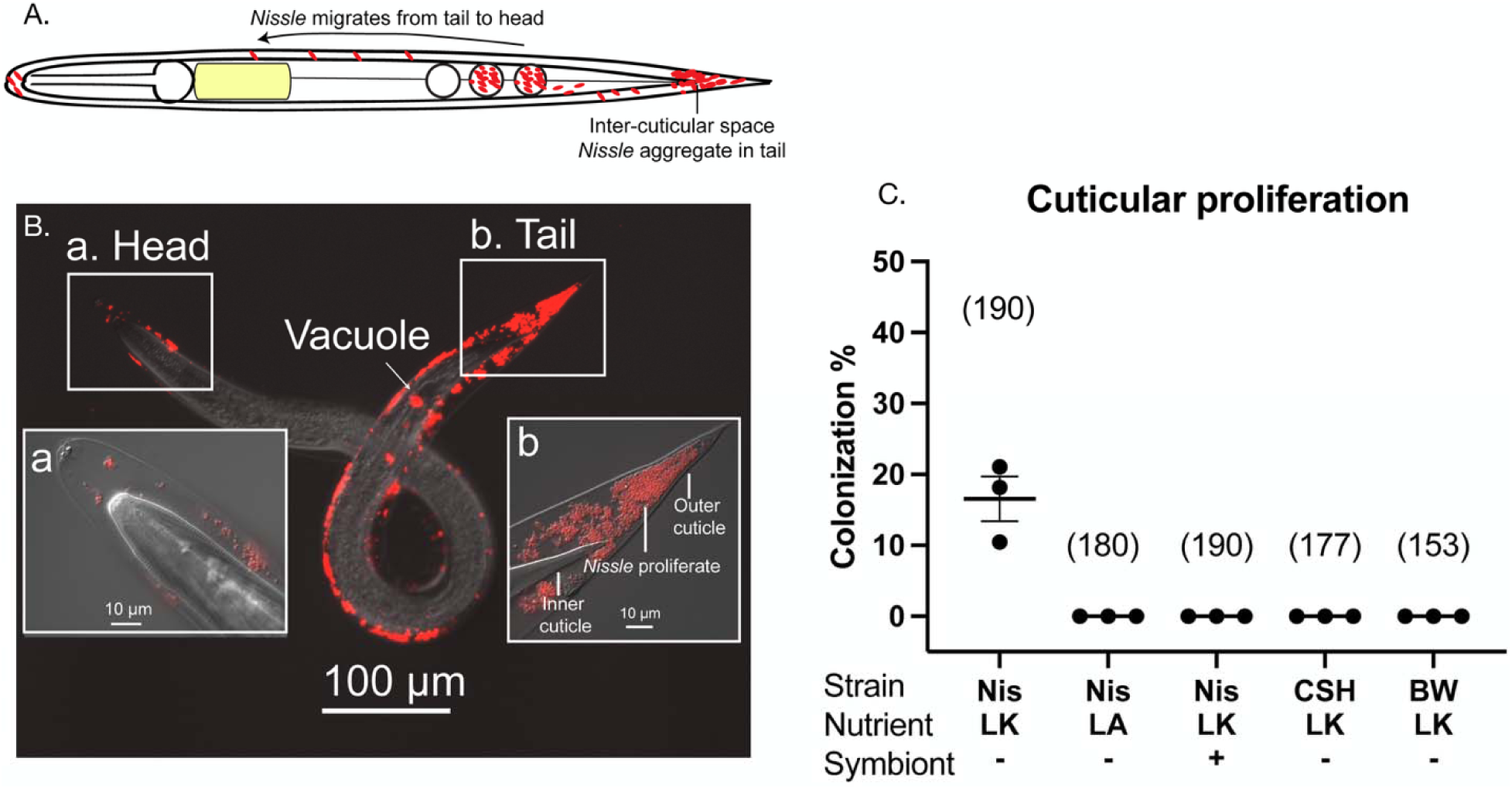
Inter-cuticular colonization of *E. coli* Nissle. (A). A schematic diagram showing *E. coli* Nissle colonizes in the inter-cuticular space in the IJs. (B). A representative IJ showing inter-cuticular proliferation of bacteria: the head region (a) and the tail region (b). Arrow indicates the intestinal vacuole colonization observed in the same IJ with cuticular colonization. (C). Quantification of cuticular proliferation frequency of *E. coli* strains: Nissle (Nis), *E. coli* CSH50 (CSH), BW25113 (BW) under nutritional conditions of either liver-kidney (LK) or lipid agar (LA). Treatments were either with (+) or without (-) symbiont *X. griffiniae.* Numbers in the parenthesis indicates the number of animals screened in each treatment.

### Proteins from *E. coli* cell lysates persist in the IJ inter-cuticular space

We observed live EcN cells that were temporarily maintained in the IJ cuticular space for approximately two weeks before they were observed to lyse. Lysis is marked by the presence of both individual bacterial cells and a smear of bright fluorescence from the mScarlet-I reporter (Fig. 5A). Eventually, no individual living mScarlet-I-tagged bacteria are observed in the intercuticular space, suggesting all the observed fluorescence comes from residual mScarlet-I proteins persisting in the bacterial lysate (Fig. 5B). Surprisingly, diffused mScarlet-I fluorescence is observed in the inter-cuticular in IJs grown under all conditions (Fig. 5C), including those where cuticular proliferation of living bacteria was not observed (Fig. 4C). This observation suggests that in addition to living bacterial colonization as described above, other mechanisms could cause bacterial proteins to be expelled and trapped in the inter-cuticular space.

**Figure 5:**
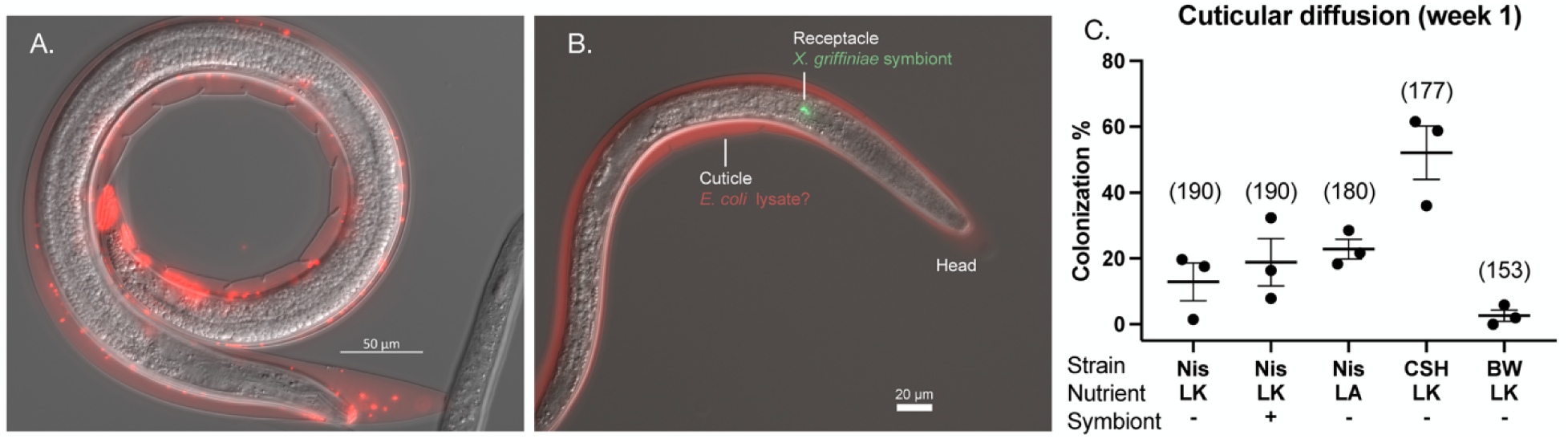
*E. coli* Nissle cells lyse in the IJ inter-cuticular space. (A). A representative IJ (2 weeks post trapping) showing transitional stage in which bacterial cells and mScarlet-I contained in cell lysate co-localize in the inter-cuticular space. (B). Diffused fluorescence from mScarlet-I expressing *E. coli* cells is found in symbiont-colonized IJs. (C). Quantification of cuticular diffusion of mScarlet-I expressing *E. coli* strains: Nissle (Nis), *E. coli* CSH50 (CSH), BW25113 (BW) under nutritional conditions of either liver-kidney (LK) or lipid agar (LA). Treatments were either with (+) or without (-) symbiont *X. griffiniae.* Numbers in the parenthesis indicates the number of animals screened in each treatment.

To test the robustness of this lysate’s fluorescence to external challenges associated with the nematode’s natural environment, we investigated if fluorescent proteins contained in the EcN lysates in the nematode cuticle can persist long-term when the IJs were challenged by external soil microbes (Fig. 6; Fig. S4). Although cuticle growth and colonization percentages varied among different replicates and experiments within populations of IJs, mScarlet-I proteins in the lysates remained fluorescent in the cuticle over eight weeks of IJ aging (Fig. 6; Fig. S4). Overall, soil challenge did not significantly affect mScarlet-I persistence in the cuticle.

**Figure 6:**
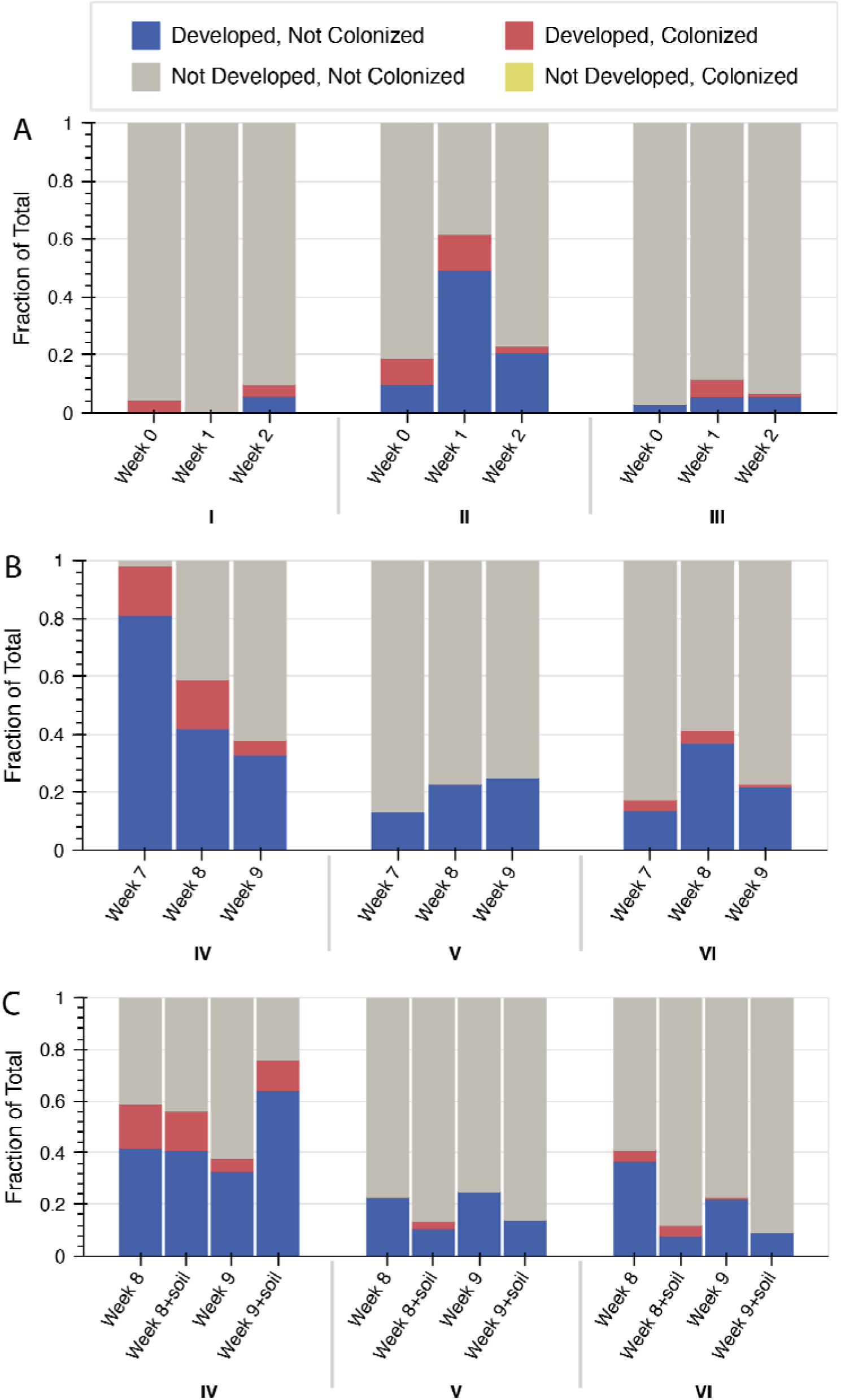
Proportion of developed outer cuticles and cuticularly colonized IJs as a fraction of total IJs with and without soil challenge. Developed cuticles (noted as ‘developed’) include those with fully developed, less developed, or broken outer cuticles. The presence of *E. coli* Nissle produced mScarlet-I proteins in IJ cuticles (noted as ‘colonized’) was assessed by fluorescence microscopy. (A). IJ cuticle development and colonization of mScarlet-I protein in the inter-cuticular space over three weeks. (I), (II), and (III) represents three biological replicates. (B). Cuticle development and cuticular mScarlet-I colonization frequency over ten weeks. (IV), (V), and (VI) represent three biological replicates. (C). A comparison of IJ cuticle development and cuticular colonization by mScarlet-I in week 8 and week 9 with and without challenge by non-sterile soil.

## Discussion

In this research, we found EcN cells interacting with four pairs of *S. hermaphroditum* coelomocytes. In model organism *C. elegans* that has the most characterized anatomy and immune system among nematode species, three pairs of coelomocytes are attached to the pseudocoelomic space with three layers of defense preventing bacterial access: the grinding of bacteria by the pharynx, multiple layers of cuticles, and an active immune system defending against the invading microbes (Altun and Hall, 2005). Here, we show that *E. coli* Nissle can access *S. hermaphroditum* coelomocytes, suggesting that at least one of the three barriers fails to defend against EcN. Invertebrate immune cells are known to interact with both mutualistic and pathogenic symbionts. For instance, in the Hawaiian bobtail squid *Euprymna scolopes,* symbiotic *Vibrio fischeri* bacteria adhere to a single blood-cell type, the haemocytes, with less affinity in comparison to pathogen adherence (Nyholm *et al*., 2009). In *S. hermaphroditum*, the mutualistic symbiont *X. griffiniae* has not been observed to interact with host coelomocytes, suggesting that the role of these immune cells may be specific to pathogens.

EcN cells were also found enclosed in the posterior intestinal pockets of IJs. These pockets, which we term ‘vacuoles’, are shown in both conventional IJs (those caught from nature and grown only through the natural insect infection process) and axenic IJs (those that are gnotobiotic). This suggests that neither the presence of EcN nor the absence of *Xenorhabdus* symbionts causes the formation of these structures. Based on the size and morphology of the debris present in the intestinal vacuoles, these void spaces are likely to have formed via cell death of the intestinal cells or organelles (Fig. 3B, panel a; Fig. S1; Video S2 and S3). Our data suggest that the posterior end, similar to the anterior end of the IJ intestine (Chaston *et al*., 2013), features compartments that are specific to the colonization and proliferation of certain environmental microbes. In previous studies, environmental microbes were found in *S. scapterisci* in the ‘inter-cuticular space’, defined as the space in between the inner cuticle of J2 and J3 larvae (Bonifassi *et al*., 1999) . Those rod-shaped bacteria, reported as ‘contaminants’, were thought to be wrapped into the J3 cuticle during nematode growth and molting (Fig. 7A). Note that in this research, EcN cells are localized to the space in between the collagenous inner cuticle and the outer cuticle. Based on our observations, we propose that the bacteria enter the inter-cuticular space after being expelled from the intestine (Fig. 4A and 4B). Within the inter-cuticular space, EcN cells were observed to migrate from the posterior to the anterior side where they form aggregates, proving that these bacterial cells are indeed living. What causes the lysis of EcN cells in the cuticle is unknown, some possibilities include host nematode immune response, or an exhaustion of nutrients in the inter-cuticular space.

**Figure 7:**
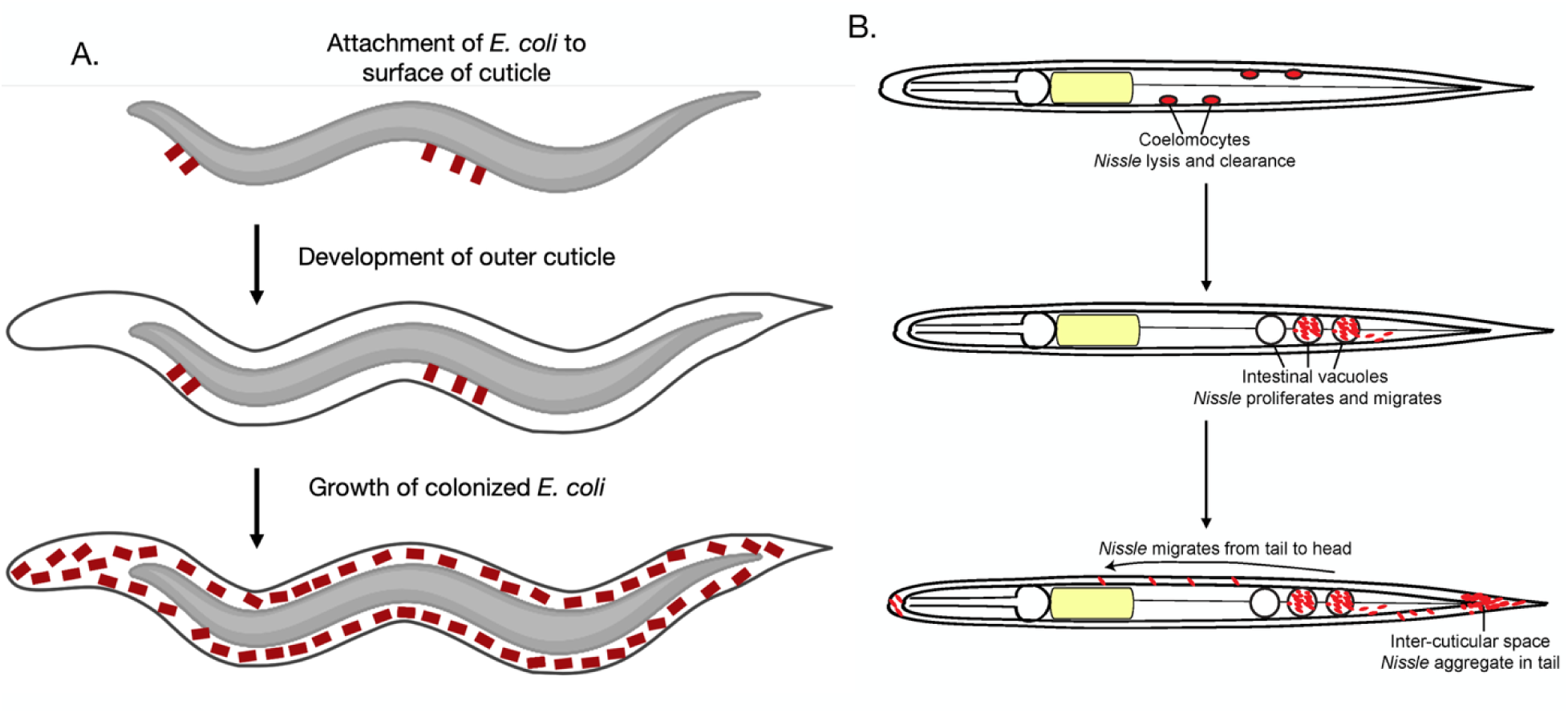
Hypothetical models of *E. coli* Nissle interaction in *S. hermaphroditum*. (A). a hypothetical model based on a previous publication of *S. scapterisci* cuticular microbes (Bonifassi *et al*., 1999) : environmental microbes attaching to the inner cuticle during IJ formation. (B). Current hypothetical model based on data from this research: *E. coli* Nissle possibly causes mild infection in *S. hermaphroditum*.

Despite the potential of binary symbiosis systems in environmental engineering, they remain underexplored. Engineering vertically transmitted symbionts or endosymbionts, which reside in host cell cytosol and are maternally passed through oocytes (Bright and Bulgheresi, 2010), is technically challenging. In contrast, horizontally transmitted microbiomes—such as the association of *S. hermaphroditum* with *Xenorhabdus griffiniae* and EcN—are acquired extracellularly from the environment in each generation. This mode of symbiont acquisition offers key advantages for microbial bioengineering: symbiotic bacteria (including mutualistic and pathogenic) can be cultured and genetically modified *ex vivo*, then easily re-associated with the host. This enables the use of engineered microbes to monitor microenvironments within host tissues or to assess external environmental conditions within relevant ecosystems.

## Conclusion

In this work, we report previously unidentified cells and tissues within *Steinernema* nematodes where *E. coli* Nissle (EcN) can localize, proliferate, and lyse. We characterized sequential EcN localization within infective juveniles (IJs), including immune cells (coelomocytes), posterior intestinal vacuoles, and the inter-cuticular space between the interior collagenous cuticle and the exterior cuticle (Fig. 7B). Remarkably, mScarlet-I proteins from EcN lysates persisted within the inter-cuticular space for over eight weeks under non-sterile soil conditions. This observation challenges the paradigm that environmental microbes are passively “wrapped” within molting nematode cuticles (Fig. 7A). Instead, our findings demonstrate that EcN can actively associate with soil-dwelling *Steinernema* in a tissue-specific manner in a niche that is separate from the nematode’s natural symbiont. During EcN invasion, the immune response of *Steinernema hermaphroditum* engages the pseudocoelom, intestinal structures, and cuticle. Based on our data in this research, we propose EcN to be a mild pathogen of *S. hermaphroditum,* which opens a new avenue of establishing *S. hermaphroditum* as a pathogenesis model. EcN cells and bacterial proteins have the potential for engineering biocircuits to monitor host-microbe interactions and soil ecosystem.

## Acknowledgements

We thank Tara Chari for sharing the *E. coli* Nissle strain with constitutive mScarlet-I reporter used in this research. We thank Heidi Goodrich-Blair, Jennifer Heppert, and Omar Alani for sharing the *Xenorhabdus* bacterial strains used in this research. We thank Paul Sternberg for providing space and equipment support.

## Supplemental Materials

**Video S1**: bacterial lysis within the *S. hermaphroditum* coelomocytes. *E. coli* Nissle expressing mScarlet is localized in the coelomocytes.

**Video S2**: intestinal cell or organelle debris in a representative intestinal vacuole.

**Video S3**: a representative IJ with two empty intestinal vacuoles housing nematode organelle debris, and two vacuoles housing EcN cells expressing mScarlet-I.

---

**Figure S1:**
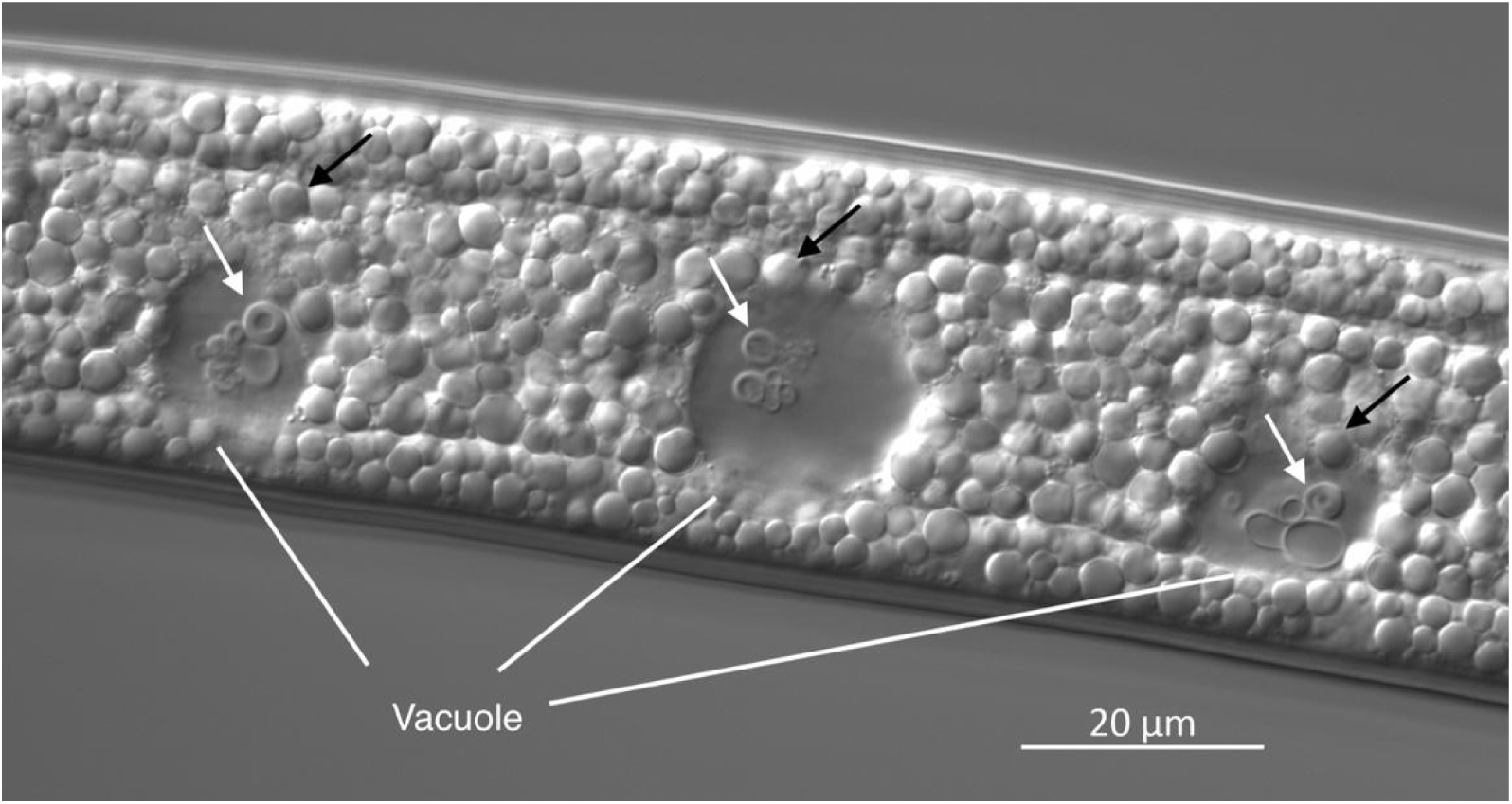
cell or organelle debris in the posterior intestinal vacuoles in a representative infective juvenile. Black arrows: gut granules; white arrows: cell or organelle debris in the intestinal vacuoles. Scale bar = 20 μm.

**Figure S2:**
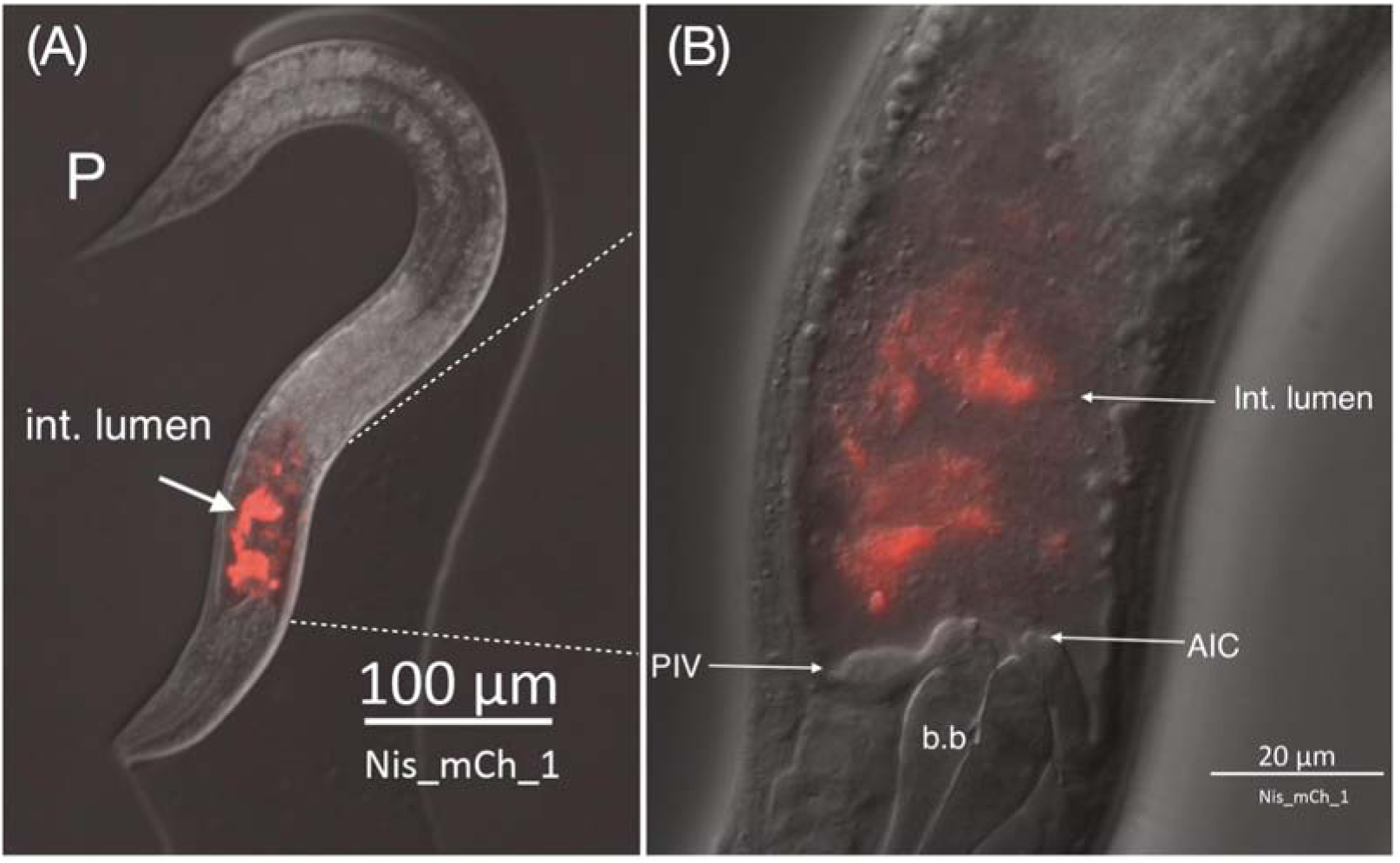
A representative *S. hermaphroditum* juvenile fed on *E. coli* Nissle. mScarlet-I expressing EcN cells are engulfed and digested in the intestinal lumen of J2 stage of *S. hermaphroditum.* ‘P’ denotes posterior end of the nematode; ‘int. lumen’ denotes ‘intestinal lumen’; ‘b.b’ denotes ‘basal bulb’; ‘AIC’ denotes ‘anterior intestinal caecum’; ‘PIV’ denotes ‘pharyngeal intestinal valve’.

**Figure S3:**
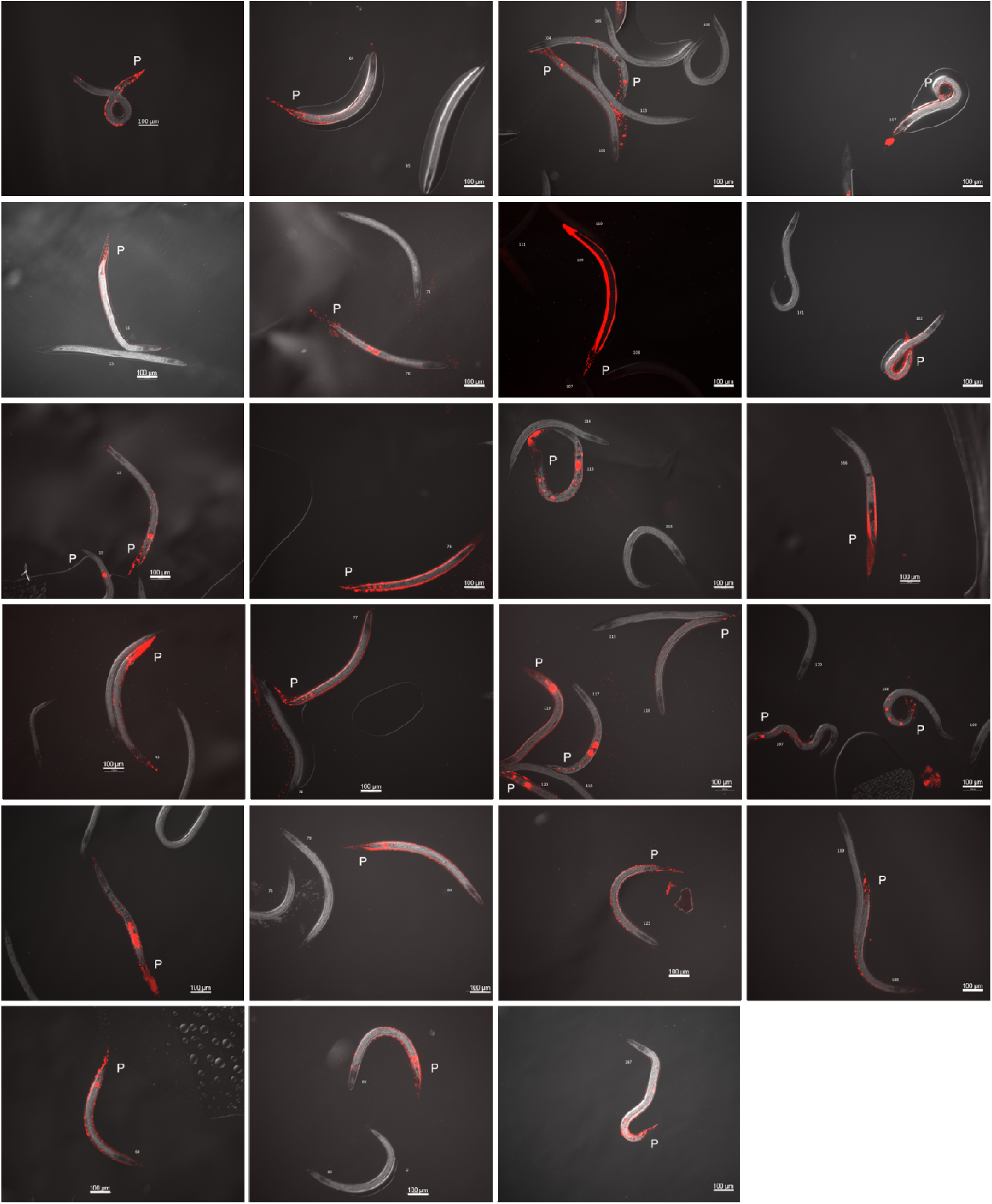
A profile of twenty-nine IJs with EcN colonization in the intestinal vacuoles and the inter-cuticualr space. ‘P’ denotes the posterior side of the IJ.

**Figure S4:**
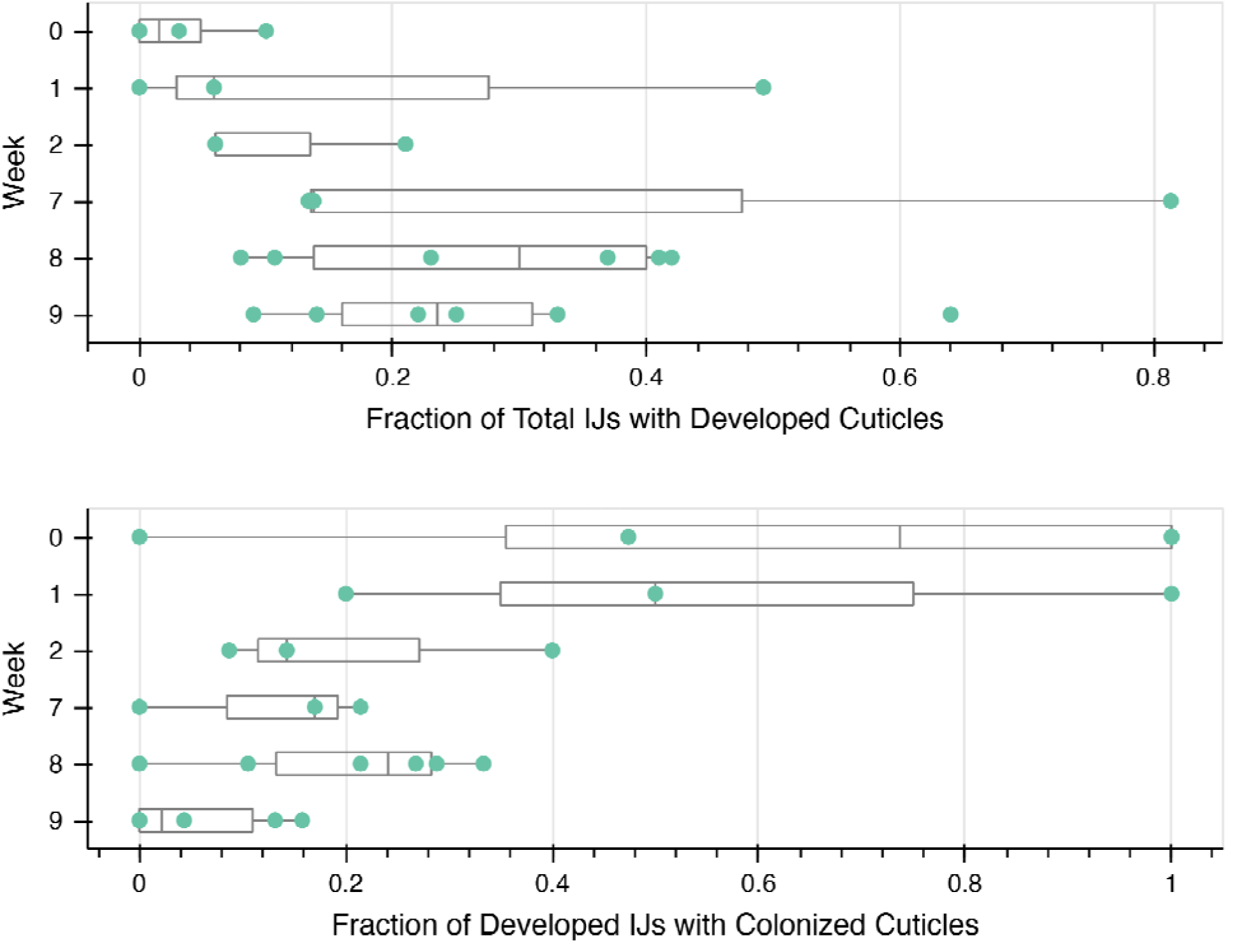
outer cuticle growth and bacterial protein (mScarlet-I) colonization in *Steinernema hermaphroditum* IJs. (A): Fractions of IJs with developed outer cuticle over 9 week’s time course of experiment. (B): Fraction of developed IJs with outer cuticle that has cuticular colonization of mScarlet from EcN lysate over the 9 weeks time course of experiment.

